# CASTpFold: Computed Atlas of Surface Topography of the universe of protein Folds

**DOI:** 10.1101/2024.05.04.592496

**Authors:** Bowei Ye, Wei Tian, Boshen Wang, Jie Liang

## Abstract

Geometric and topological properties of protein structures, including surface pockets, interior cavities, and cross channels, are of fundamental importance for proteins to carry out their functions. Computed Atlas of Surface Topography of proteins (CASTp) is a widely used web server for locating, delineating, and measuring these geometric and topological properties of protein structures. Recent developments in AI-based protein structure prediction such as AlphaFold2 (AF2) have significantly expanded our knowledge on protein structures. Here we present CASTpFold, a continuation of CASTp that provides accurate and comprehensive identifications and quantifications of protein topography. It now provides (i) results on an expanded database of proteins, including the Protein Data Bank (PDB) and non-singleton representative structures of AlphaFold2 structures, covering 183 million AF2 structures; (ii) functional pockets prediction with corresponding Gene Ontology (GO) terms or Enzyme Commission (EC) numbers for AF2-predicted structures; and (iii) pocket similarity search function for surface and protein-protein interface pockets. The CASTpFold web server is freely accessible at https://cfold.bme.uic.edu/castpfold/.

## 1 Introduction

Protein structures are complex, containing a multitude of surface pockets, internal cavities, and interconnected channels. These distinctive topographical and topological features provide the micro-environments essential for the biochemical functions of the proteins, such as binding with ligands, interacting with DNA, and catalyzing enzymatic reactions. Identification and measurement of these topographical features are of fundamental importance in deciphering the relationship between protein structures and functions (1), in studying protein fitness (2; 3), and in developing therapeutic interventions (4).

The CASTp server provides detailed quantitative analysis of protein topographical and topological features (5–7) and is widely used in various applications, including exploring therapeutics for neuropsychiatric disorders (8), drugging “undruggable” pockets (9), elucidating G-protein coupling specificity (10), unraveling lipid translocation mechanisms in autophagy (11), understanding bacteriophage host ranges (12), and investigating plant metabolism processes (13).

Recently, the advent of advanced AI tools such as AlphaFold2 (AF2) (14) has significantly expanded the repository of available protein structures. The AlphaFold Protein Structure Database (AFDB) provides more than 214 million predicted protein structures. As AF2- predictions provide valuable resources of protein structures, here we present the CASTpFold server, which retains all critical functionalities of its predecessors while providing topographical and topological quantification to the AF2-predicted structures, so that the research community could have a comprehensive tool for analyzing and understanding the spatial arrangement, function, and similarity of protein topography in both predicted and PDB protein structures, facilitating more informed investigations.

## 2 MATERIALS AND METHODS

### 2.1 The CASTpFold server

The CASTpFold server is based on the alpha shape method (15) from topological data analysis to identify topographical and topological features and to compute their areas, volumes, and imprints (16–20). Topographical concave regions of proteins include surface pockets and depressions, CASTpFold considers pockets, which are on the protein surface and are accessible through a narrow entrance from the outside openings large enough for a probe ball, e.g., a water molecule, to access (Fig 1A-➀). Pockets differ from depressions, which are concave surface areas without entrance constriction (Fig 1A-➂). As an example, Fig 1B depicts a pocket in an AF2- predicted structure, which is predicted to be a functional region involved in kinase activity. Cavities are topological features and are internal voids buried inside a protein that the probe ball cannot access (Fig 1A-➁). Channels are a class of pockets with two mouth openings at opposite sides. An example is shown in Fig 1C (PDB ID: 5BUN, Pocket ID: 1). CASTpFold identifies and measures surface pockets and interior cavities on a protein structure, as well as protein-protein interface (PPI) pockets and cavities where multiple proteins or subunits interact to form protein complexes.

**Figure 1.**
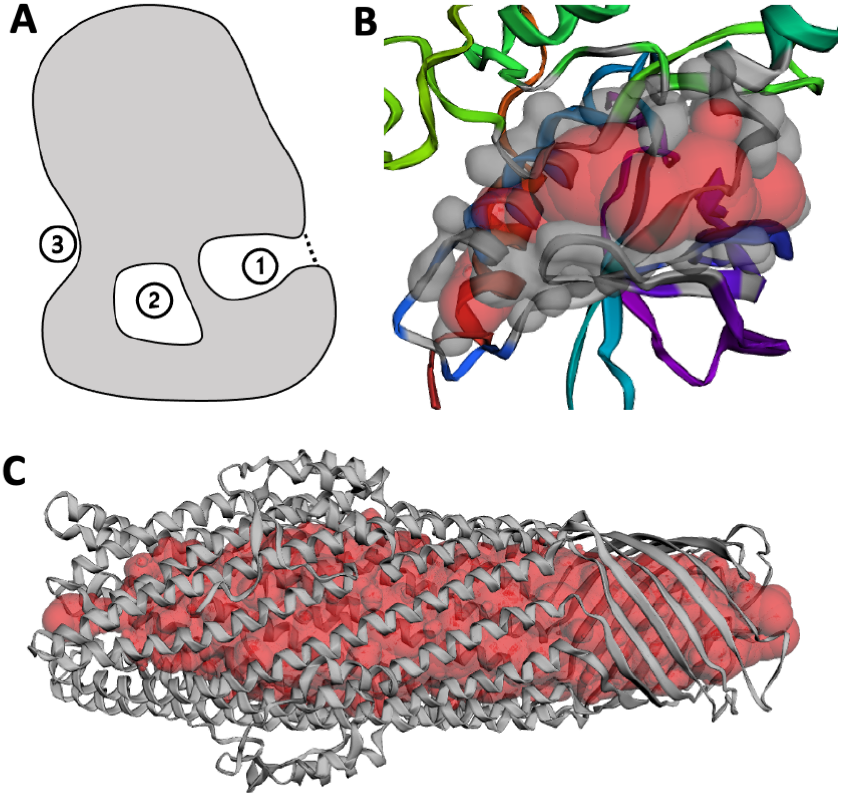
The concave regions identified on protein structures. (A) Protein surfaces contain three types of concave regions: ➀ an accessible surface pocket with a narrowed entrance, indicated by a dashed line; ➁ a completely enclosed cavity (void) lacking an entrance; and ➂ a shallow depression. (B) A surface pocket supported by atoms in grey as identified by CASTpFold, which exhibits kinase activity (GO:0016301, Pocket ID: 2, style: “surface”) in the AF2-predicted structure (AF2 ID: A0A2N0QT79). Its imprint is in red. (C) A cross channel in the outer membrane protein ST50 with its imprint highlighted in red balls (PDB ID: 5BUN, Pocket ID: 1).

The CASTpFold server also includes other information about the protein. The secondary structures are assigned using the DSSP method (21). The protein residue annotations are obtained from the UniProt database (22) and aligned with the PDB structure via the SIFTS database (23). Functional pockets for AF2 structures identified by CASTpFold are annotated with GO terms or EC numbers by integrating results obtained from the deep learning algorithm DeepFRI (24; 25). The pocket similarity is measured using an approach adapted from the Foldseek method (26).

### 2.2 New features of CASTpFold

#### 2.2.1 Enlarged database

CASTpFold expands its scope beyond the PDB database (27) and now includes all non-singleton AF2 representative structures (28), providing an exhaustive analysis of protein topography of most of the protein-universe. Specifically, the 214 million AF2-predicted structures (as of Nov 2022) can be grouped into 18,661,407 structural clusters, by the criterion that their representative structures are recognized by Foldseek (26). For protein sequences in the UniProt database, after removing those labeled as “fragments”, the remaining 2,302,908 non-singleton sequences clusters (as of Dec 2023) can be mapped to the 183,581,108 AF2-predicted structures (28), whose representative structures are all included in the CASTpFold server. When querying any of the 183,581,108 AF2-predicted structures, the search is automatically directed to its appropriate representative AF2 structure. For example, a query for the structure (AF2 ID: R4G0B4) redirects to its representative structure (AF2 ID: A0A6B1EM21), ensuring users can efficiently find the most relevant structural information.

#### 2.2.2 Predicted functional pocket for AF2 structures

The UniProt database (22), with its repository of over 250 million sequences, has only about 0.3% of its entries manually reviewed in UniProtKB/Swiss-Prot. Given that protein functionalities are enabled by specific local surface regions, pinpointing relevant topographical features such as pockets and cavities can help decipher the mechanisms of protein functions. To identify potential functional pockets and cavities in unannotated AF2-predicted structures, we combine protein topography analysis with functional residue identification obtained through Deep- FRI (24; 25), so our computed functional pockets along with DeepFRI derived GO terms or EC numbers are provided. An example is shown In Fig 2A, where two surface pockets (Pocket IDs: 1 in lime and 2 in red) in an AF2-predicted structure (AF2 ID: A0A0C3Q0M4) are predicted to facilitate kinase activity. Here the computed negative imprint of the pocket volume are colored lime and red, respectively. Fig 2B illustrates a member of a PPI pocket cluster (cluster size: 294), highlighted in red, located at the functional interface between the SARS-CoV-2 Spike protein and the human angiotensin-converting enzyme-2 (ACE2) protein (PDB ID:6M0J, Pocket ID: 1).

**Figure 2.**
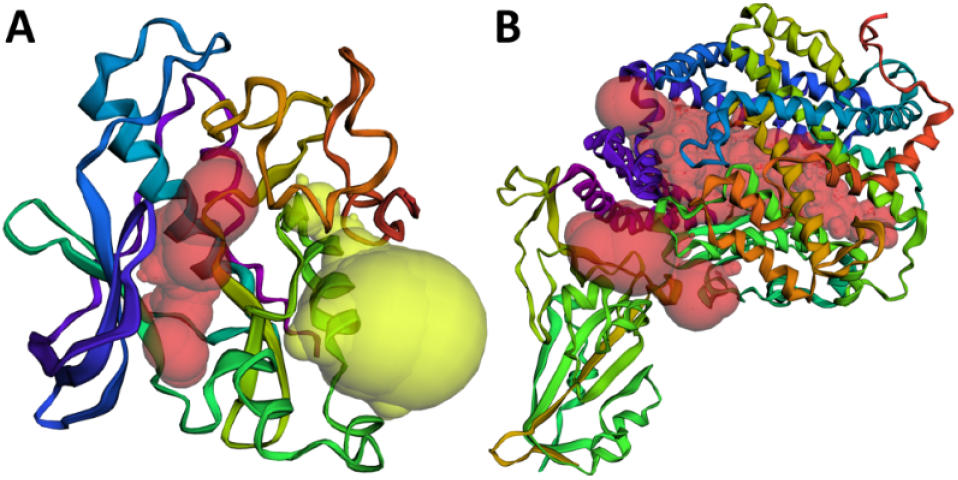
Surface pockets and protein-protein interface pockets: (A) Two surface pockets identified on the AF2-predicted structure (AF2 ID: A0A0C3Q0M4, Pocket IDs: 1 in lime and 2 in red). They are collectively predicted to carry out kinase activity; (B) A member of a PPI pocket cluster (cluster size: 294), located at the interface of the SARS-CoV-2 Spike/ACE2 Complex (PDB ID: 6M0J, Pocket ID: 1)

#### 2.2.3 Pocket similarity search

In our dataset of PDB structures and 2.3 million AF2-predicted representative structures, we have collected 4,108,408 surface pockets and 433,078 protein-protein interface (PPI) pockets, each satisfying the requirement of containing a minimum of 14 residues. The surface and PPI pocket databases have been clustered using the Foldseek algorithm (26) for rapid identification of similar pockets. Users can explore the relationship among the surface and PPI pockets by clicking the “Pocket Similarity” panel, where they can access, download, and visualize lists of similar pockets.

## 3 INPUT AND OUTPUT

### 3.1 Input

The CASTpFold server processes protein structures in PDB/mmcif format, incorporating a user-specified probe radius for detailed topographic analysis of the solvent-accessible surface. Users can either explore pre-computed results by searching with standard PDB/AF2 IDs through the server’s interface, or submit their own protein structures for computation. Pre-computed results were derived using a default probe radius of 1.4 Å for water. For customized computation requests, users have the option to adjust the probe radius within the range of 0 to 10 Å, enabling customized analysis.

### 3.2 Output

The CASTpFold server identifies all surface pockets, interior cavities, and cross channels in a protein structure, offering precise mapping of every atom involved in these topographic features. Additionally, it quantifies their exact volumes and surface areas, including the dimensions of any mouth openings. These calculations are performed analytically through the solvent-accessible surface model (SA) (29) and the molecular surface model (MS) (30). The “Pocket Info” panel will display only SA-related data, while MS-related information is available in the downloadable content, including the absolute values and ratios of atom-level surface exposed areas and volumes for both SA and MS models. User submitted structures will be subject to detailed topographical analyses, the results of which can be accessed and downloaded via CASTpFold. Predictions of functional pockets and measurements of pocket similarities are also accessible, although they are currently restricted to the precomputed structures.

### 3.3 Case study

The predicted functional pocket shown in Fig 1B is further illustrated in Fig 3, which presents a pocket related to potential kinase activity identified in the AF2-predicted structure (AF2 ID: A0A2N0QT79, Pocket ID: 2).

**Figure 3.**
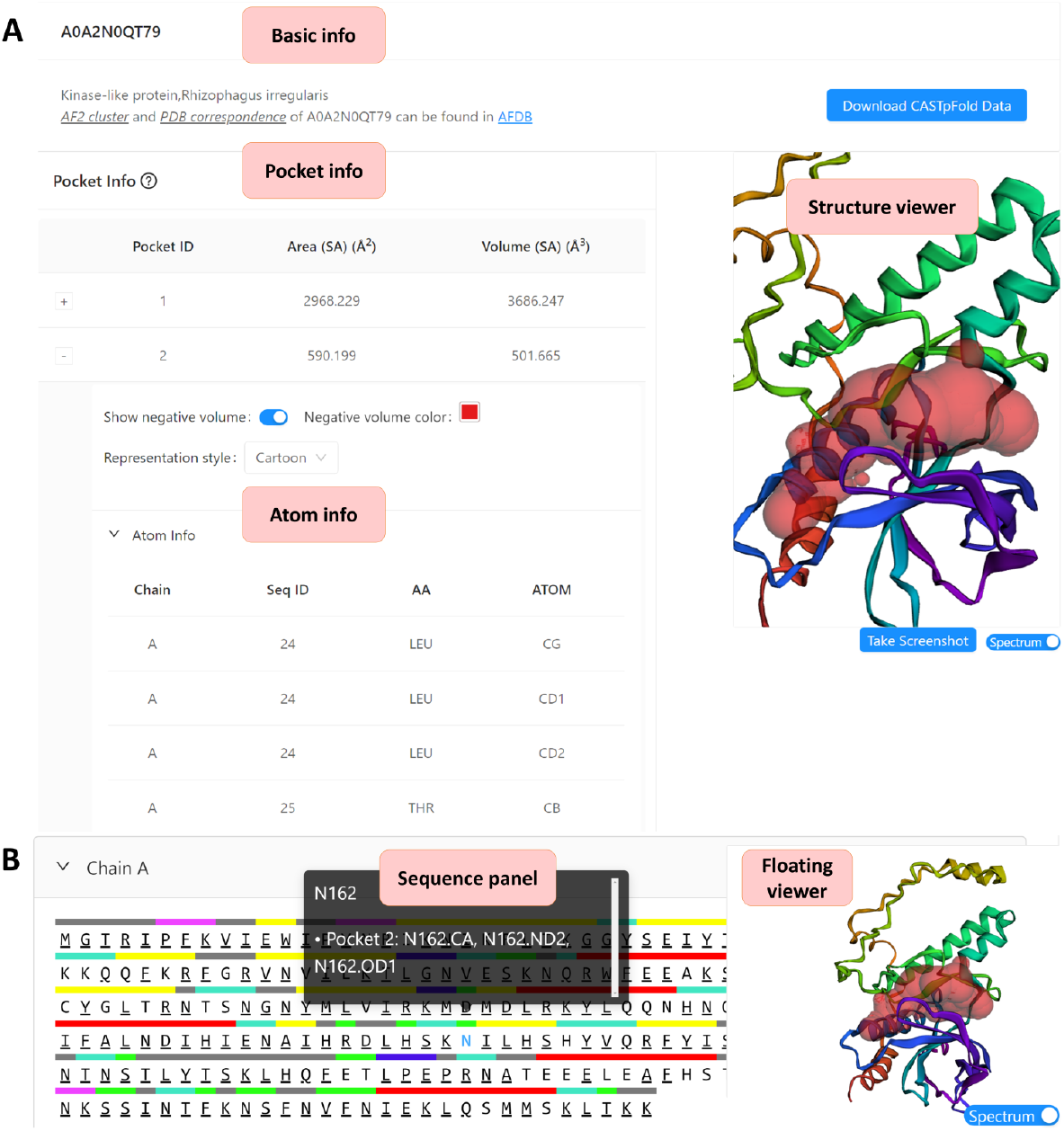
The main user interface of the CASTpFold server supports both precomputed and user-uploaded structures. (A) The server displays data on pocket surface area, volume, and detailed information on atoms contributing to the pocket. Here the second pocket is highlighted in “cartoon” style. (B) The sequence panel presents information on atomic contributions and annotations of residues along the sequence. Here the residue “N162” and its atoms contributing to the pocket formation are listed(Pocket ID:2). In addition, a structure viewer and a floating viewer facilitate visualization of the results.

In Fig 3A, the “Basic Info” panel for protein (AF2 ID: A0A2N0QT79) displays information from the UniProt database. The “Pocket Info” panel depicts the selected pocket (Pocket ID: 2), which is rendered in cartoon style in the “Structure viewer” panel. Additionally, an expandable “Atom Info” panel reveals the atomic details of the selected pocket. Fig 3B shows the “Sequence Info” panel for the structure, including pocket residue with atom information and annotations. The viewer panel is designed to float, maintaining visual access while navigating the site, and users can save the current view by using the “Take Screenshot” button.

While navigating the CASTpFold server, users exploring AF2-predicted structures can find a “Predicted Function” panel, which lists the predicted function of the pocket (Fig 4A), including the associated GO terms or EC numbers, functionally relevant positions, and links to details of potential functions. For PDB structures, an “Annotation” panel is available (Fig 4B), providing annotations extracted from the UniProt database (22). AF2 predicted functionally relevant residues or PDB annotated residues can be visualized together with their respective pockets in a combined view (Fig 4A and 4B). Additionally, a “Pocket Similarity” panel is accessible for both AF2 and PDB structures, allowing users to discover pockets similar to their query protein across other proteins in the database. For example, Fig 4C shows a pocket from an AF2- predicted structure (AF2 ID: A0A2N0QT79, Pocket ID: 2) that has a total of 93 pockets across both PDB and AF2 structures exhibiting similarity to the queried pocket. One of these is from the PDB structure (PDB ID: 7NQQ, Pocket ID: 1), which is visualized in the “Similar pocket viewer”. The “PDB counterpart in AFDB” panel is designed for PDB structures to display their AF2 counterparts, enabling users to compare PDB structures with their AF2 equivalents by clicking the corresponding links (Fig 4D).

**Figure 4.**
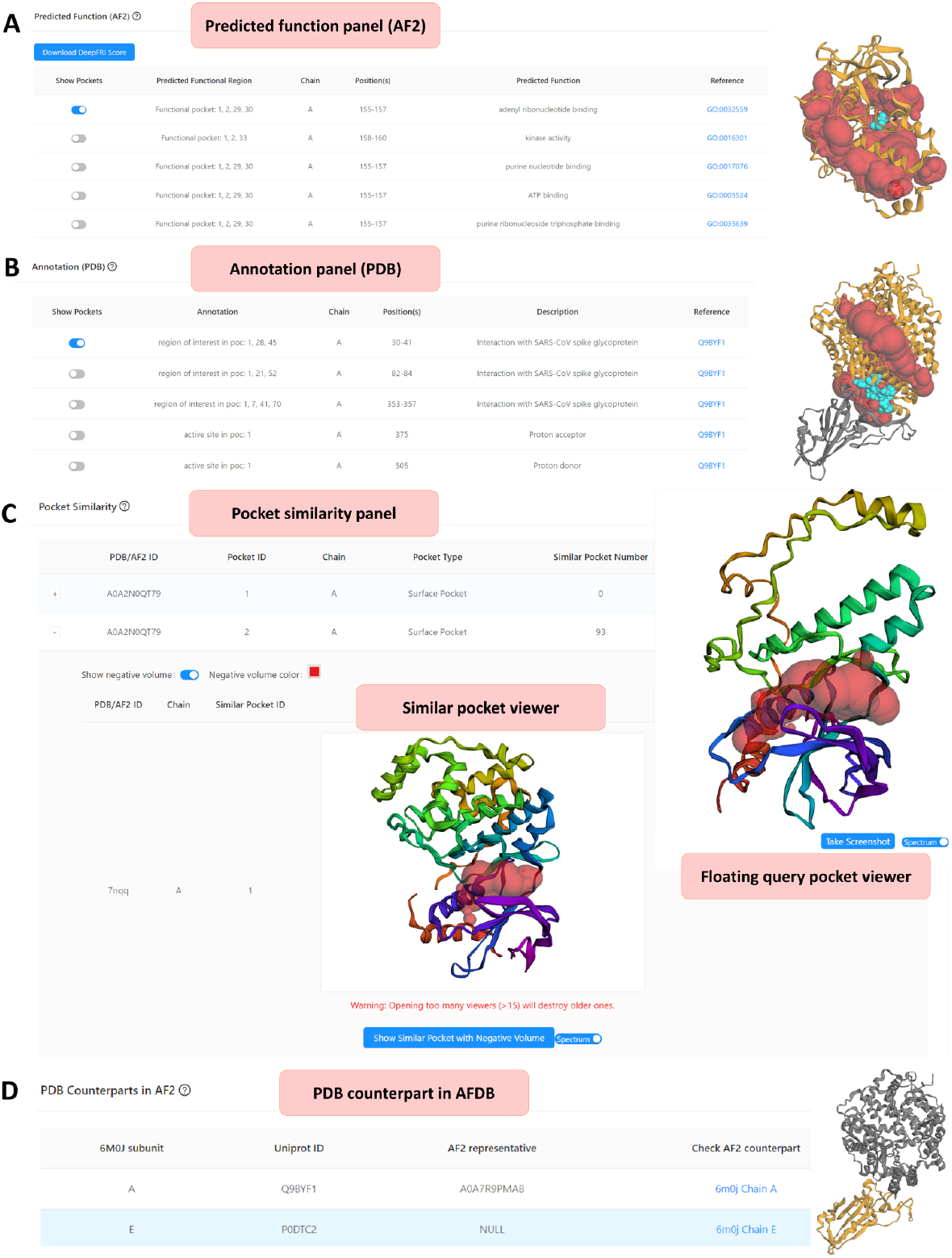
Additional features of the user interface of the CASTpFold server for precomputed structures. (A) The predicted function panel displays functional pockets in AF2 structures with links to relevant GO terms or EC numbers. The predicted functionally relevant positions are shown as cyan balls, their corresponding chains in orange, and the imprints of the pockets in red for combined viewing. (B) The annotation panel displays UniProt-sourced annotations for PDB structures, highlighting annotated residues, their respective chains, and their relevant pocket imprints in cyan, orange, and red, respectively. (C) The pocket similarity panel offers a list of pockets from other proteins similar to the query pocket, for both AF2 and PDB structures. (D) The “PDB counterpart in AFDB” panel shows the AF2 counterparts of all subunits of the query PDB structure, where the corresponding Uniport ID is mapped to its AF2 representative entry. Users can perform a detailed comparison by exploring the topographies of the AF2 counterpart by opening the counterpart link.

## 4 DISCUSSION

CASTpFold represents a major update of the CASTp server, with the following significant enhancements: (i) an expanded database that includes PDB entries and 2.3 million AF2 structures representing 183 million AF2 structures, covering over 85% of the AFDB; (ii) predicted functional pockets of AF2 structures, complete with associated GO terms or EC numbers; (iii) a pocket similarity search function of surface and PPI pockets for both PDB and AF2 structures; (iv) the user interface has been redesigned to be more informative, ensuring improved user engagement. These advancements are expected to further facilitate the exploration and analysis of protein structures and their functions. CASTpFold is free and open to all users and there is no login requirement.

## 5 FUNDING

The grant support of NIH R35GM127084 is gratefully acknowledged.

### 5.0.1 Conflict of interest statement

None declared.

## Notes

https://cfold.bme.uic.edu/castpfold/

### Competing Interest Statement

The authors have declared no competing interest.

